# Changes in life history and population size can explain relative neutral diversity levels on X and autosomes in extant human populations

**DOI:** 10.1101/763524

**Authors:** Guy Amster, David A. Murphy, William M. Milligan, Guy Sella

## Abstract

In human populations, relative levels of neutral polymorphism on the X and autosomes differ markedly from each other and from the naive theoretical expectation of ¾. These differences have attracted considerable attention, with studies highlighting several potential causes, including male biased mutation and reproductive variance, historical changes in population size, and selection at linked loci. We revisit this question in light of our new theory about the effects of life history and given pedigree-based estimates of the dependence of human mutation rates on sex and age. We demonstrate that life history effects, particularly higher generation times in males than females, likely had multiple effects on human X-to-autosomes (X:A) polymorphism ratios, through the extent of male mutation bias, the equilibrium X:A ratios of effective population sizes, and differential responses to changes in population size. We also show that the standard approach of using divergence between species to correct for the male bias in mutation results in biased estimates of X:A effective population size ratios. We obtain alternative estimates using pedigree-based estimates of the male mutation bias, which reveal X:A ratios of effective population sizes to be considerably greater than previously appreciated. We then show that the joint effects of historical changes in life history and population size can explain X:A polymorphism ratios in extant human populations. Our results suggest that ancestral human populations were highly polygynous; that non-African populations experienced a substantial reduction in polygyny and/or increase in male-biased generation times around the out of Africa bottleneck; and that extant diversity levels were affected by fairly recent changes in sex-specific life history.

**Significance Statement:** All else being equal, the ratio of diversity levels on X and autosomes at selectively neutral sites should mirror the ratio of their numbers in the population and thus equal ¾. In reality, the ratios observed across human populations differ markedly from ¾ and from each other. Because from a population perspective, autosomes spend an equal number of generations in both sexes while the X spends twice as many generations in females, these departures from the naïve expectations likely reflect differences between male and female life histories and their effects on mutation processes. Indeed, we show that the ratios observed across human populations can be explained by demographic history, assuming plausible, sex-specific mutation rates, generation times and reproductive variances.

## Introduction

Neutral polymorphism patterns on the X and autosomes reflect a combination of evolutionary forces. Everything else being equal, the X to autosome (X:A) polymorphism ratio should be ¾, because the number of X-chromosomes in a population is ¾ that of autosomes. A complication, however, is that autosomes spend an equal number of generations in diploid form in both sexes, whereas the X spends twice as many generations in diploid form in females as in haploid form in males. As a result, the X:A polymorphism ratio can also be shaped by differences in male and female life history and mutation processes, as well as by differences in the effects of demographic history and selection at linked sites on the X and autosomes. The effects of these factors have been studied theoretically (1) and in relation to observations in many species (2–7). Notably, their effects on polymorphism ratios in human populations has garnered considerable interest over the past decade (8–15).

The impact of selection at linked sites on neutral diversity levels could differ for X and autosomes because of differences in recombination rates, in the density of selected regions, and in the efficacy and modes of selection. Notably, the hemizygosity of the X in males leads to a more rapid fixation of recessive or partially recessive beneficial alleles and to a more rapid purging of recessive deleterious ones (16, 17). Accounting for these effects and for recombination rates suggests that in humans—in mammals more generally—the effects of selection at linked sites should be stronger on the X ((18), but see (2)). To evaluate these effects empirically, several studies have examined how polymorphism levels on the X and autosomes vary with genetic distance from putatively selected regions, e.g., from coding and conserved non-coding regions (10, 12, 14, 19–21). In most hominids, including humans, such comparisons confirm the theoretical expectation that selection at linked loci reduces X:A ratios (10, 20, 21). They further suggest that the effects are minimal sufficiently far from genes (10, 19), thereby providing an opportunity to examine the effects of other factors shaping X:A ratios in isolation, by considering regions that are minimally affected.

Even far from genes, however, the X:A ratios in humans and other hominids differ markedly from the naive expectation (10, 12, 14). Polymorphism levels on the X and autosomes are typically divided by divergence from an outgroup (e.g., divergence to orangutan or rhesus macaque is used to normalize polymorphism levels in humans) in order to control for the effects of higher mutation rates in males and variation in mutation rates along the genome (8). The normalized estimates of X:A ratios in regions far from genes range between ¾ and 1 among human populations, generally decreasing with the distance from Africa (10, 12, 21, 22). Ratios exceeding ¾ have also been observed in most other hominids ((14), but see (15)).

These departures from ¾ and differences among populations and species have been attributed in part to the effects of demographic history, in particular to historical changes in population size. If we assume that the effective population size on the X is generally smaller than on autosomes, then changes in population size will have a different impact on polymorphism levels on X and autosomes (4, 23–25). Notably, population bottlenecks that occurred sufficiently recently, such as the Out of Africa (OoA) bottleneck in human evolution, will have decreased the X:A ratio, because a greater proportion of X-linked lineages will have coalesced during the bottleneck (25). Indeed, simulation studies suggested that historical changes in population size have contributed substantially to the X:A ratios decrease with the distance from Africa (21). Historical differences between males and females may have also played a role. For example, Keinan and colleagues speculated that male biased migration or longer male generation times during the Out-of-Africa bottleneck contributed to the lower X:A ratios in non-Africans (11).

Sex differences in life history traits are also likely to have had substantial effects on X:A ratios. The most straightforward of these effects arises from higher reproductive variances in males than in females (e.g., due to sexual selection (8)), which cause higher coalescence rates on autosomes and thus increased X:A ratios (1, 8). This increase is theoretically bound by a multiplicative factor of 3/2 (1), but is probably much smaller in reality. Nonetheless, greater male reproductive variances in extant hunter-gatherers and hominid species (26), suggest that male biased variance plausibly contributed to observed differences in X:A ratios, as well as to their departure from ¾.

In addition, higher generation times in males may have also had substantial yet underappreciated effects (27). Higher generation times in males decrease coalescence rates on autosomes compared to the X and thus the X:A ratio of effective population sizes (27). In addition, mutation rates in humans–likely in mammals more generally–increase more rapidly with paternal than with maternal age (28–33). Longer generation times in males therefore decrease mutation rates on the X relative to autosomes. Normalizing polymorphism estimates by divergence to an outgroup may not fully account for this mutational effect if, as is likely, male mutation bias evolve over phylogenetic time scales (34–36); moreover, normalized ratios also reflect a non-mutational generation times effect on X:A divergence ratios, as longer generation times in males imply fewer generations on autosomes relative to the X since the species split (36, 37). Thus, male and female generation times can in principle affect X:A ratios in multiple ways, which should be considered jointly.

Here we examine these effects, and those of life history more generally, on polymorphism ratios in humans. We begin with general considerations: about the effects in populations of constant size, about the effects in response to changes in population size, and about biases introduced by normalizing polymorphism ratios by divergence to an outgroup. We then estimate X:A polymorphism ratios in six human populations in which historical changes in population size were inferred previously, and show that considering these effects jointly can explain the observed ratios.

## Results

### Life history effects in populations of constant size

In a parallel paper (27), we derive expressions for neutral X:A polymorphism ratios in a panmictic population of constant size, under a model that captures quite general life history effects. The model assumes that the population is divided into sex specific age classes, with female and male proportions *γ*_*F*_ and *γ*_*M*_ at birth, respectively (*γ*_*F*_ + *γ*_*M*_ = 1), and that the sizes of subsequent age classes of each sex declines with age, reflecting sex and age specific mortality. Fecundity also depends on sex and age and incorporates sex-specific reproductive variances and correlations in the numbers of offspring at different ages. Generation times in females, *G*_*F*_, and in males, *G*_*M*_, are defined as the expectations of maternal and paternal ages. Mutation rates can vary with sex and age, with their per generation rates in females, *μ*_*F*_, and in males, *μ*_*M*_, defined as expectations over parental ages. The expected numbers of offspring of each sex necessarily equals 1, but female and male reproductive variances, *V*_*F*_ and *V*_*M*_ respectively, may differ due to sex and age dependent mortality and fecundity.

We show that the X:A ratio of effective population sizes is then:

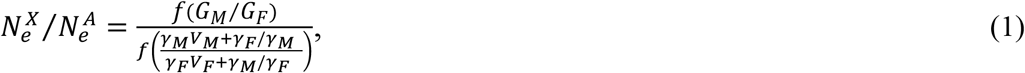

where 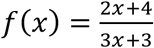. Here we define the effective population sizes such that they equal the number of individuals under the standard Wright-Fisher model, but we note that they are sometimes defined as the inverse of coalescence rates (e.g., in the statement that all else being equal, the X:A ratio of *N*_*e*_ is ¾). We refer to the X:A ratio of inverse coalescence rates as the genealogical ratio, which in this case is simply ¾ times the ratio of *N*_*e*_ values.

We also show that the X:A ratio of expected heterozygosities is

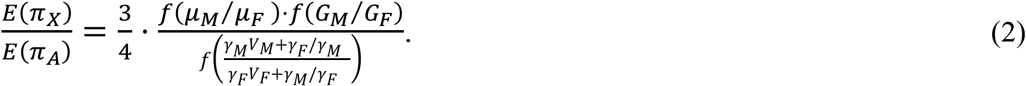

When the mutation rates, generation times, numbers of newborns, and reproductive variance are identical in both sexes, the heterozygosities ratio and the genealogical ratio reduce to the naïve neutral expectation of ¾. When these factors differ between sexes, Eq. 2 provides a simple expression for the effect of each factor. Notably, the effects of sex specific age and reproductive structure reduce to the effects of male to female ratios of mutation rates (*α* = *μ*_*M*_/*μ*_*F*_), generation times (*G*_*M*_/*G*_*F*_), and terms combining reproductive variances and birth proportions ((*γ*_*M*_*V*_*M*_ + *γ*_*F*_/*γ*_*M*_)/(*γ*_*F*_*V*_*F*_ + *γ*_*M*_/*γ*_*F*_)). As the proportions of females and males at birth in humans are nearly equal, we henceforth assume that they are, and thus that the latter ratio equals (2 + *V*_*M*_)/(2 + *V*_*F*_); for brevity, we refer to it as the ratio of reproductive variances.

All three ratios in Eq. 2 are male-biased in many taxa (38–40). We examine their effects on X:A polymorphism ratios in humans, given available estimates. Sex-specific reproductive variances were measured in five extant hunter-gatherer groups, albeit using small sample sizes, and found to be 1.7-4.2 folds higher in males (26), with reproductive variances ratios corresponding to a 6-20% increase in X:A polymorphism ratios. Sex-specific generation times were measured in seven hunter-gatherer groups, with mean generation times found to vary between 25 and 33 years and generation times ratios between 1.03 and 1.37 (36, 38), corresponding to a 0.5%-5.2% decrease in X:A polymorphism ratios. Male mutation bias, *α*, was estimated in pedigree studies (32), and found to increase approximately linearly on the sex-ratio of generation times, *G*_*M*_/*G*_*F*_ (33), and depend negligibly on the sex-average of generations times ((33); Fig. 3). Based on these estimates, male mutation bias would decrease X:A polymorphism ratios by ~18% even without male biased generation times, and by ~21% with *G*_*M*_/*G*_*F*_ = 1.37 (the upper end estimated in extant hunter gatherers). Fig. 1 shows the joint effects of the ratios of reproductive variances and generation times (including both genealogical and mutational effects) on human X:A polymorphism ratios, assuming a constant population size.

**Figure 1.**
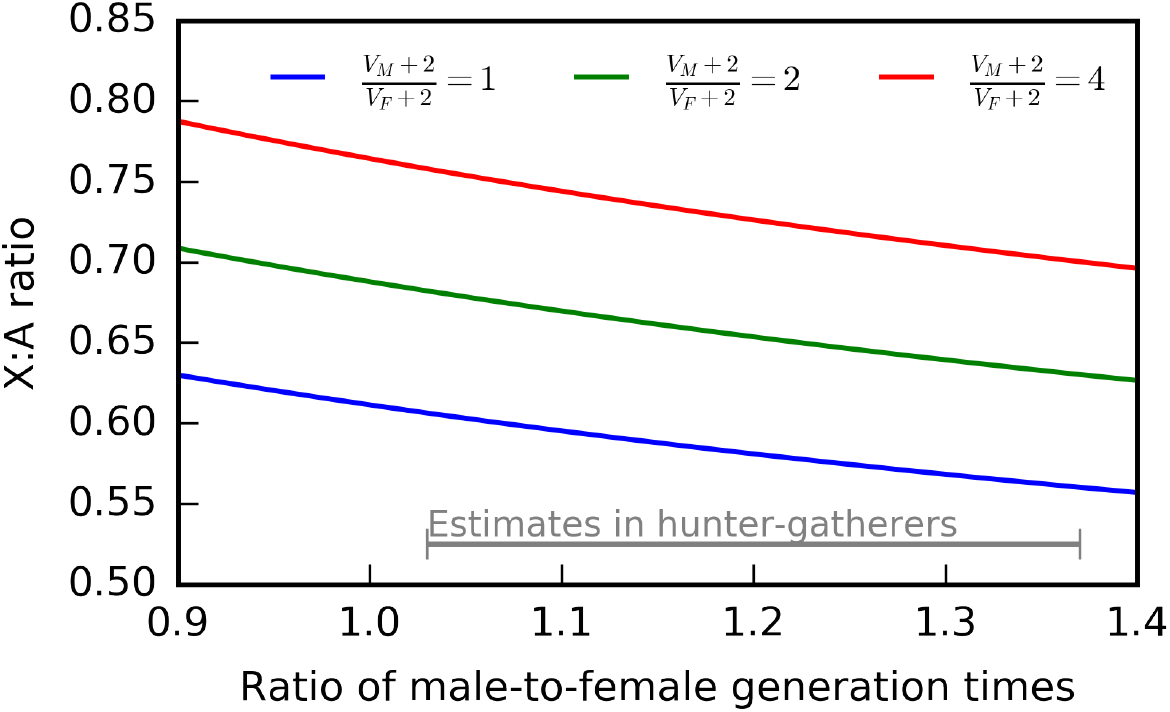
Life history and mutational effects on the X:A ratio assuming a constant population size. Note that these predictions are not directly comparable with most ratios reported in the literature, which are normalization by divergence (see below).

### Life history effects in populations of changing size

Changes in population size affect X:A ratios (25), in ways that are modulated by sex-specific life history (27). In Amster and Sella (27) we derive closed forms for these effects, and here, as illustration, we consider a simple bottleneck scenario roughly corresponding to that of the Out of Africa (OoA) bottleneck in humans (Fig. 2). In the absence of sex differences in life history, with an equilibrium X:A genealogical ratio of ¾, diversity levels on the X experience more of a reduction during, and a faster recovery after, the bottleneck, because both periods are longer in coalescence units on the X (25). For OoA bottleneck parameters, these effects lead to a reduction in the polymorphism ratio (Fig. 2: black curve). Higher reproductive variance in males increases the equilibrium genealogical ratio, moving it closer to 1, leading to a smaller reduction in the polymorphism ratio in response to the bottleneck (Fig. 2B: blue curve). In contrast, a longer generation time in males results in a large reduction to the X:A polymorphism ratio, for two reasons (Fig. 2: red curve). It decreases the equilibrium genealogical ratio, which accelerates the response to the bottleneck on the X relative to the autosomes *in generations*, and it also decreases the generation time on the X relative to autosomes, which increases the relative number of generations on the X *per unit time* (in years). More generally, when life histories are sex specific, they modulate the effects of changes in population size on X:A polymorphism ratios in two ways: they influence the coalescence time scale of the response of X vs autosomes *in generations* and they change the generation times on X vs autosomes and thus the relative number of generations *per unit time* (in years). These consideration clarify that even if we had perfect knowledge of effective population sizes of the X and autosomes throughout history, let alone only of autosomes, we would still have to know past values of life history traits in order to predict the effects of demography on the X:A polymorphism ratio (Figs. 2 and S1).

**Figure 2.**
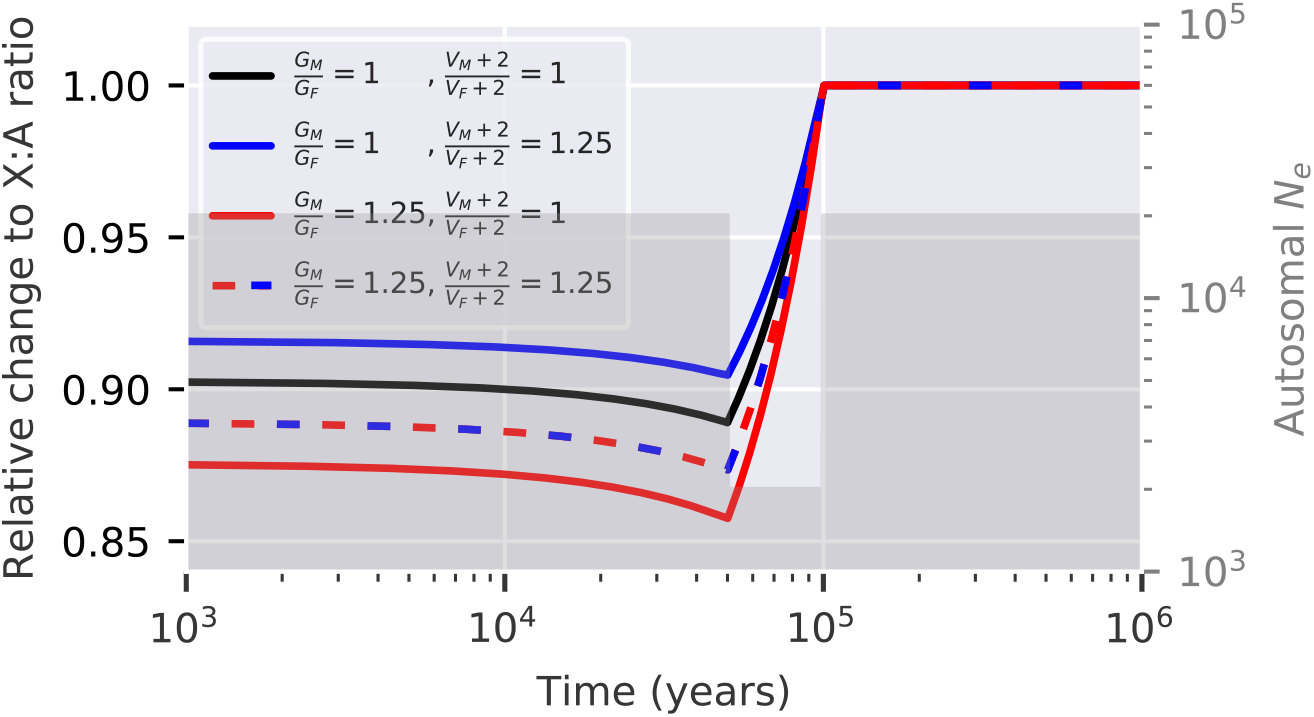
The effect of a population bottleneck on the X:A polymorphism ratio is modulated by sex-specific life history. Bottleneck parameters were chosen to roughly correspond with autosomal estimates for the OoA bottleneck in humans (right y-axis). We consider the relative change in X:A ratios (left y-axis) assuming an autosomal generation time of 30 years, and four combinations of generation times and reproductive variances ratios. When both ratios have the same value then the equilibrium genealogical ratio is ¾ (black and dashed blue-red curves); when *G*_*M*_/*G*_*F*_ = 1 then *G*_*X*_/*G*_*A*_ = 1, and when *G*_*M*_/*G*_*F*_ = 1.25 then *G*_*X*_/*G*_*A*_ = 0.72. See Fig. S1 for the absolute rather than relative changes in polymorphism and genealogical ratios.

### Normalizing ratios of polymorphism using divergence

Most studies estimate X:A genealogical ratios by dividing polymorphism levels by estimates of the number of substitutions since the split from an outgroup (e.g., (8, 10, 12, 41)). This “normalization” is meant to control for differences in mutation rates on the X and autosomes, due to male mutation bias and differences in base composition. We now ask whether this practice is valid, and in particular whether it controls for male-biased mutation when we account for life history effects on polymorphism and substitution rates.

To this end, we rely on Eq. 2 for the polymorphism ratio (ignoring changes in population size) and on a parallel expression for the substitution ratio

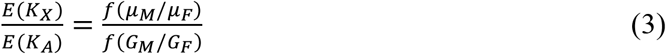

(36, 37). This allows us to derive an explicit form for the normalized ratio:

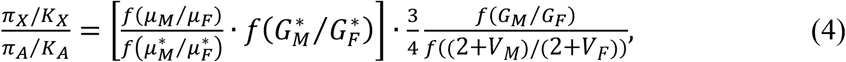

where by ‘*’ we denote parameters averaged over the lineage on which substitutions are measured (the specific form of averaging is detailed in (36)). The second term (on the right-hand side) is the X:A genealogical ratio. The first term (in brackets) includes the mutational effect on the polymorphism ratio and the terms introduced by the normalization. For the normalization to fulfill its purpose of canceling out the effect of male mutation bias, this term should equal 1. Previous work suggests that male mutation bias (*α* = *μ*_*M*_/*μ*_*F*_) evolves substantially over phylogenetic time scales (34, 42), and therefore the mutational term 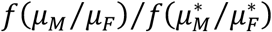 is unlikely to cancel out. However, even if it did, the dependence of the substitution ratio on the generation times ratio introduces an additional term, 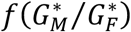. Both terms are likely to lead to bias in estimates of the genealogical X:A ratio in a given population. Furthermore, as the degree of male mutation bias probably varies among populations (e.g., due to variation in generation times (38)), relative estimates of genealogical ratios in different populations will likely biased as well.

We can assess the severity of these biases for estimates of human X:A genealogical ratios by comparing divergence- and pedigree-based estimates of the male mutation bias, *α* (Fig. 3). If differences in the mutation rate between the X and autosome arise predominantly from the male mutation bias (see Discussion), then we would expect estimates of the mutational effects on X:A polymorphism ratios based on contemporary pedigree studies to be more reliable. Indeed, while mutation rates may have evolved over the period in which neutral diversity in extant human populations arose (e.g., on the order of ~1.5 MY (43); see Discussion), such changes were likely smaller than the changes over phylogenetic time scales (e.g., on the order of ~15 and ~30 MY for divergence from orangutans and rhesus macaques, respectively (43)). Second, while both divergence and pedigree-based estimates of *α* depend on the sex ratio of generation times, in pedigree studies, this dependence reflects only the effect of generation times on mutation rates (as opposed to the non-mutational term 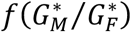 for divergence) and is explicit. Pedigree-based estimates of *α*, again assuming a linear dependence on *G*_*M*_/*G*_*F*_ (32) and estimates of *G*_*M*_/*G*_*F*_ in extant hunter-gatherers (36, 38), range between 3.6 to 4.5, and estimates in most societies point to the higher end of this range. These estimates are approximately *twofold greater* than those based on human-orangutan or human-macaque divergence (Fig. 3), commonly used to normalize human X:A ratios (8, 10, 12, 21). This strongly suggests that current estimates of human X:A genealogical ratios are substantially biased.

**Figure 3.**
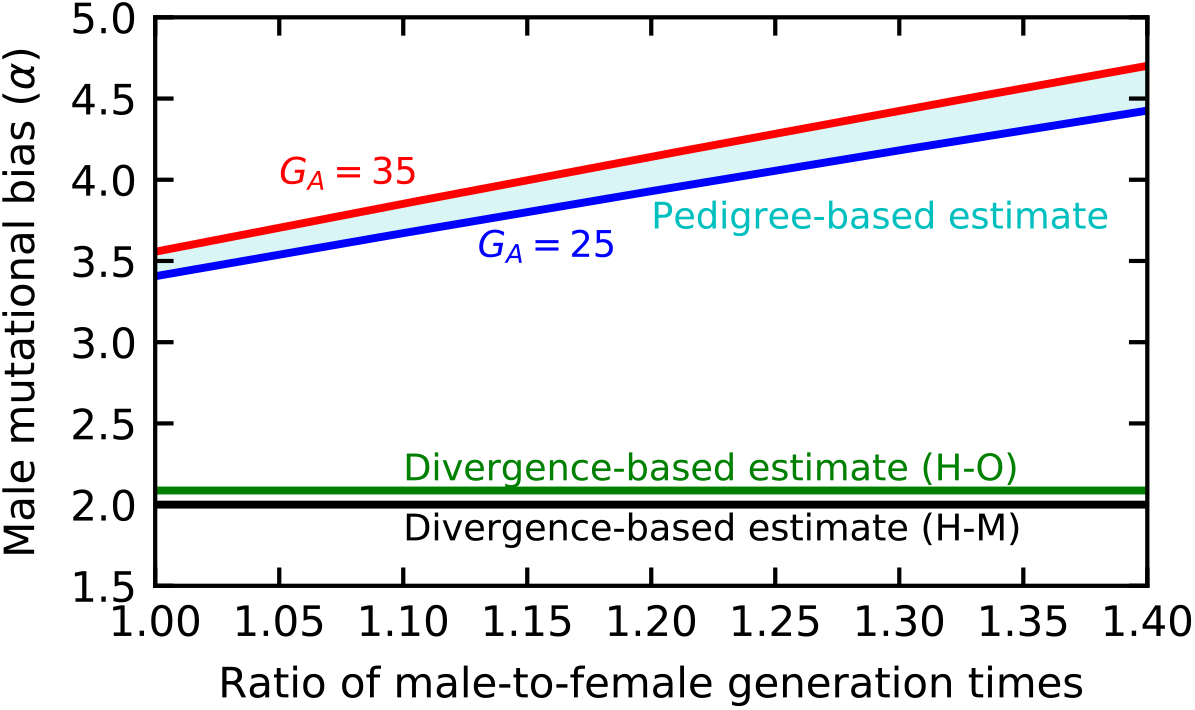
Marked difference between estimates of male mutation bias (*α*) in humans based on X:A ratios of divergence to an outgroup (SI Section 2.3 and Table S3) and on contemporary pedigree studies (SI Section 1; (32)). Pedigree-based estimates strongly depend on, and are therefore shown as a function of, the generation times ratio, *G*_*M*_/*G*_*F*_. They depend only weakly on the average generation time, *G*_*A*_, as shown by the (cyan) range corresponding to *G*_*A*_ between 25-35 years.

### Revised estimates of human X:A genealogical ratios

We therefore revisit the estimation of genealogical X:A ratios in human populations. We first estimate X:A polymorphism ratios normalized by divergence, in order to correct for local variation in mutation rates on X and autosomes (8), and then we rely on pedigree studies to correct for the bias in divergence-based estimates of *α*.

We estimate normalized, neutral polymorphism ratios in the absence of selection at linked sites in two ways (SI Section 2.2). First, we apply the standard method (8), based on measuring polymorphism and divergence at putatively neutral sites far from exons. Second, rather than imposing a threshold distance from exons, we use sites throughout the genome and rely on the McVicker et al. *B*-maps to correct for the effects of selection at linked sites (19); this approach allows us to use more data. Our estimates based on the two approaches are consistent (Fig. S5) and we henceforth rely on estimates using the second approach (Fig. 4), which are more precise. For unclear reasons, they do not agree with the estimates of Arbiza et al., which rely on similar data but are slightly higher in YRI and much higher in other populations (21).

**Figure 4.**
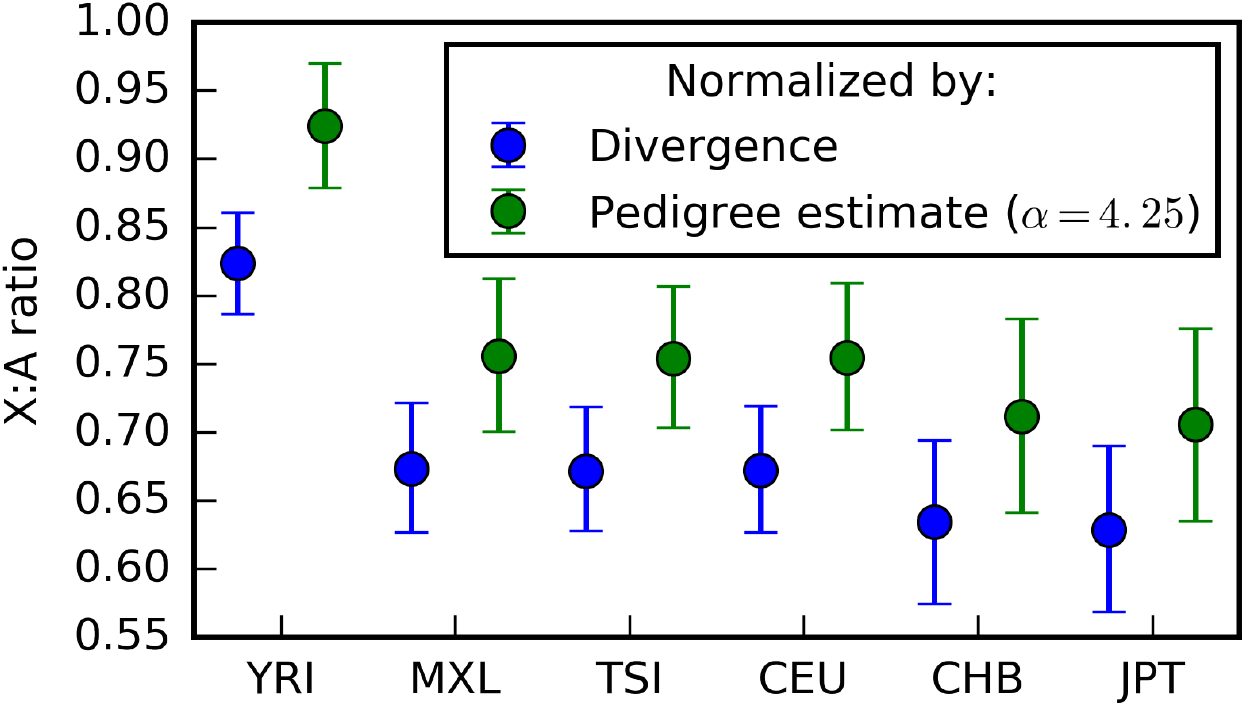
Genealogical X:A ratios in human populations are greater than previously appreciated.

To examine how much male-mutation bias affects estimates of X:A genealogical ratios, we assume *α* = 4.25, corresponding to the pedigree based estimate with the average *G*_*M*_/*G*_*F*_ measured in extant hunter-gatherers (Fig. 3). We then obtain the corrected estimates by multiplying our divergence-normalized estimates by 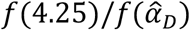, where 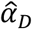 is the divergence-based estimate of male mutation bias. Dividing by 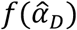 also removes the potential effect of differential levels of ancestral polymorphism on X and autosomes. The resulting estimates are *~11% greater* than those based on divergence alone, suggesting that the genealogical X:A ratios in humans are considerably greater than previously appreciated (Fig. 4).

As we already noted, pedigree-based estimates of *α* strongly depend on *G*_*M*_/*G*_*F*_. Since this ratio likely varies over time and among populations, we cannot estimate genealogical X:A ratios that reliably. This limitation is not specific to us, but instead highlights the difficulty of teasing apart the mutational and genealogical effects on X:A polymorphism ratios without making explicit or implicit assumptions about male mutation bias and its evolution.

### Explaining genealogical ratios in human populations

Instead, we turn the question on its head and ask whether the effects of sex ratios of generation times and reproductive variances as well as historical changes in population size could explain the estimates of X:A polymorphism ratios that are normalized by divergence. To this end, we rely on pairwise MSMC-based estimates of historical, autosomal effective population sizes for the six 1000G populations in which these were inferred (Fig. 7A; (44)). In all cases considered, we assume that *G*_*A*_ = 30, to match the assumptions of these previous demographic inferences (44). We first consider the ratio in YRI, then the reduction in ratio in CEU relative to YRI, and lastly, the ratios in all six populations jointly.

### Polymorphism ratio in YRI

The X:A polymorphism ratio in YRI is remarkably high (Fig. 4), which is indicative of substantial polygyny (i.e., that a minority of males sired offspring with multiple females). Accounting for historical changes in population size and assuming, for example, the average generation times ratio estimate in extant hunter-gatherers (*G*_*M*_/*G*_*F*_ =1.26), this ratio implies a reproductive variance ratio of (2 + *V*_*M*_)/(2 + *V*_*F*_) ≅ 5.5 (95% CI [3.9, 8.2]) and thus high reproductive variance in males (*V*_*M*_ ≥ 14.4). More generally, matching the estimated X:A ratio in YRI with constant life history ratios defines a trade-off in which assuming a higher generation times ratio implies more extreme male-biased reproductive variances (Fig. 5). Given that the generation times ratio was likely greater than 1, our findings suggest substantial polygyny (where a minority of males sire offspring with multiple females) in the ancestors of YRI, and thus of other human populations (8).

**Figure 5.**
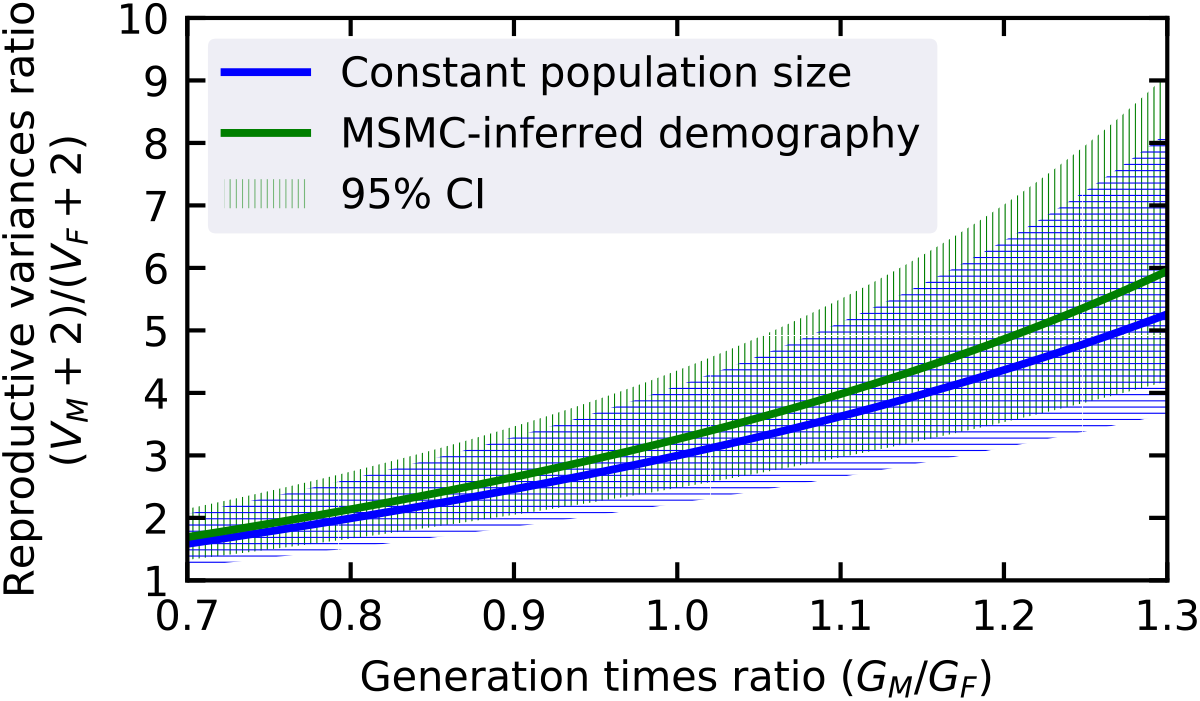
The combinations of sex ratios of generation times and reproductive variances that are consistent with estimates (± 95% CI) of the X:A polymorphism ratio in YRI.

### Reduced polymorphism ratio in CEU relative to YRI

To explain the 18.4% reduction in polymorphism ratio in CEU relative to YRI, we must assume that sex-specific life history varied over time and between populations. Without life history effects (model i in Fig. 6; (21)), historical changes in population size and the OoA bottleneck in particular lead to about half of the observed reduction. Yet sex ratios of generation times and reproductive variances in the past almost certainly differed from 1 and varied over time and among populations (e.g., as they do among extant hunter-gatherers; (38)). To explore the potential effect of such variation, we consider several models in which values of *G*_*M*_/*G*_*F*_ are constrained to be between 0.9 and 1.4 and values of (*V*_*M*_ + 2)/(*V*_*F*_ + 2) are constrained to be between 1 and 2.5; these ranges are somewhat arbitrary, yet they are clearly possible, given estimates in extant hunter-gatherers (38). Requiring the ratios to have been the same in both populations and constant over time (model ii in Fig. 6), we find that the maximal reduction in the X:A polymorphism ratio in CEU relative to YRI is 12.3% (see SI Section 3 for details on the maximization). Allowing the ratios to have different values before and after the split between YRI and CEU but requiring them to be the same in both populations (model iii in Fig. 6), results in only a slightly greater maximal reduction of 12.7% in the X:A polymorphism ratio. Further allowing for population specific parameter values after the split (model iv in Fig. 6), we find that the maximal reduction in the ratio in CEU relative to YRI rises to 20%, which is greater than the reduction observed. These results illustrate, quite surprisingly, that fairly recent changes to life history traits (relative to the average age of neutral polymorphism in either population) can dramatically affect X:A polymorphism ratios. In particular, they show that the reduction in polymorphism ratio in CEU relative to YRI can be explained by assuming that life history parameters varied within plausible ranges over time and among populations.

**Figure 6.**
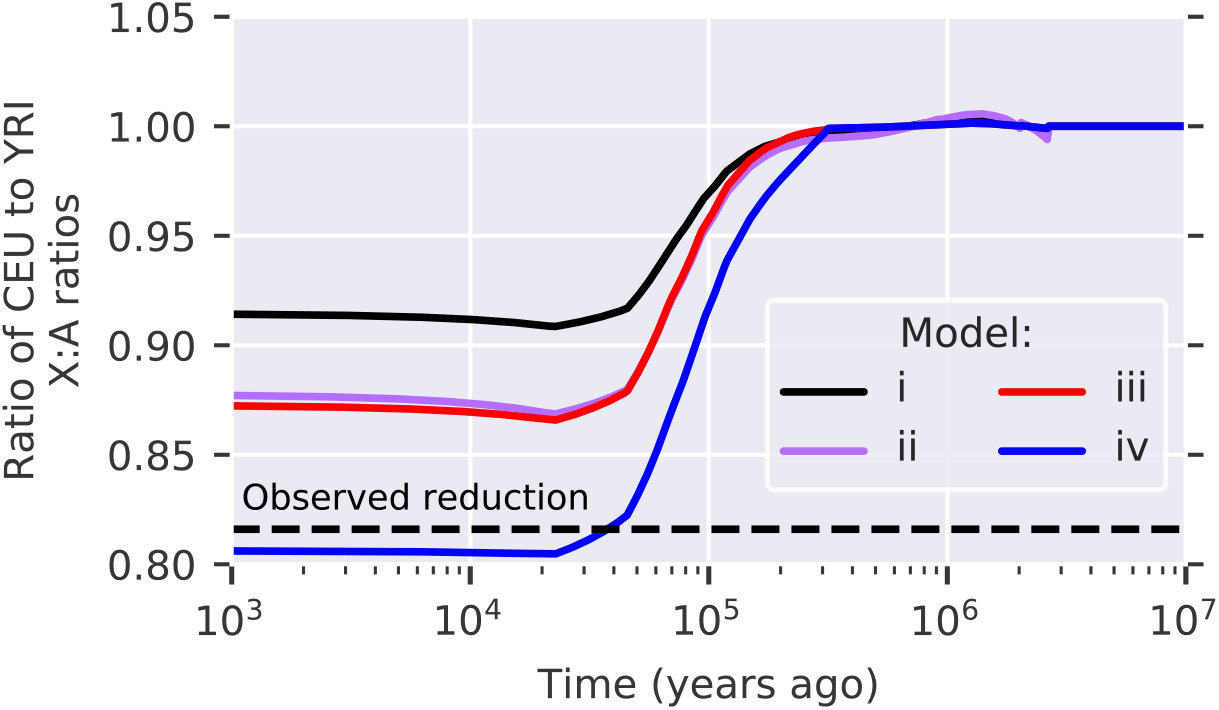
The expected relative reduction in polymorphism ratios between YRI and CEU under inferred historical changes in population size (Fig. 7A) and different models of life history (see text for details). (i) without life history effects: *G*_*M*_/*G*_*F*_ = (2 + *V*_*M*_)/(2 + *V*_*F*_) = 1. (ii-iv) The ratios (2 + *V*_*M*_)/(2 + *V*_*F*_) and *G*_*M*_/*G*_*F*_ are allowed to vary within the ranges detailed in the text, and are chosen to maximize the extant reduction in polymorphism ratios in CEU relative to YRI under the following constraints: (ii) constant ratios over time and populations; (iii) ratios can differ before and after the populations split but are the same in both populations; (iv) ratios are the same before the populations split but different after. The estimated reduction in polymorphism ratio in CEU relative to YRI is shown for comparison.

**Figure 7.**
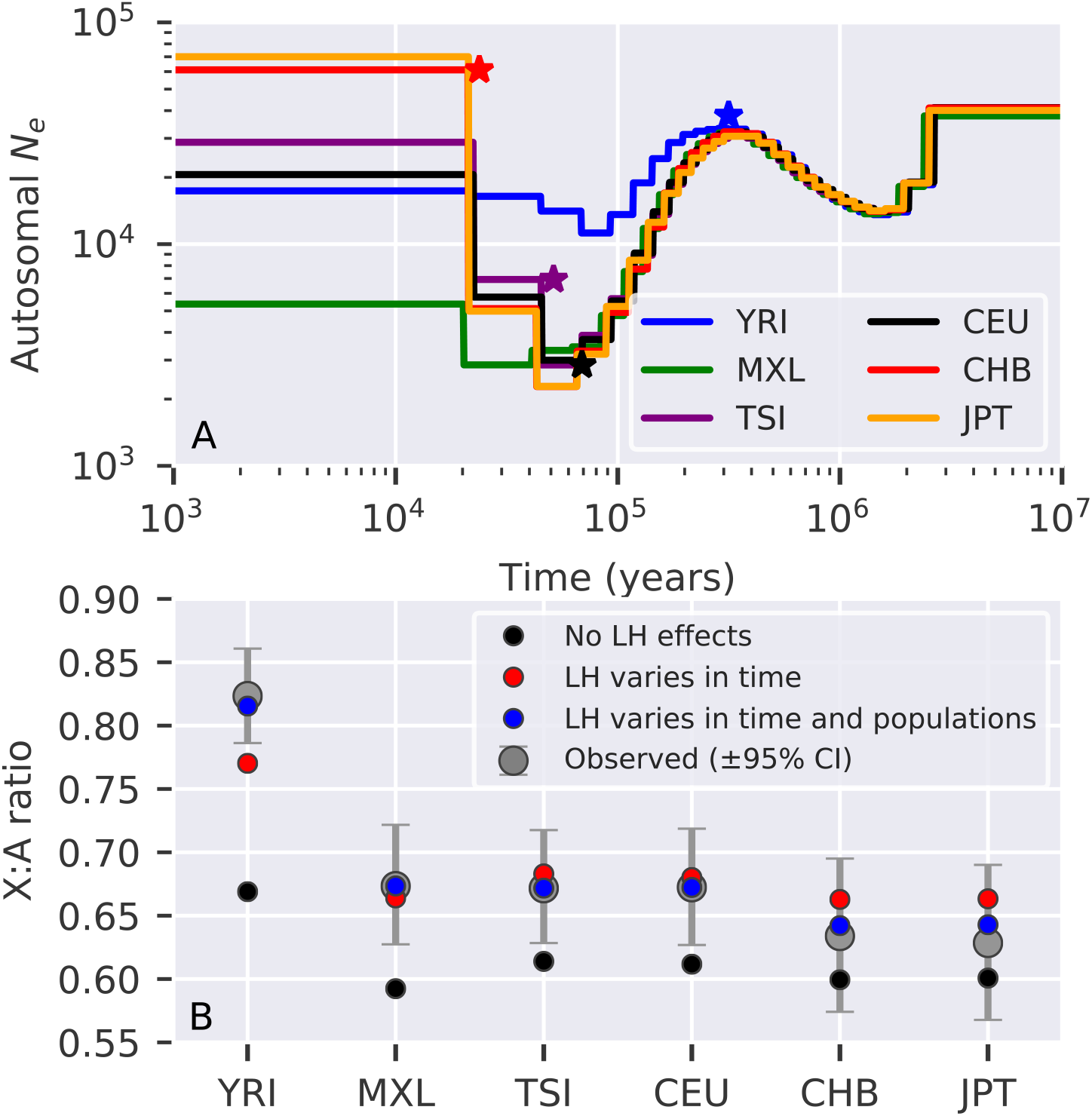
Variation in sex-specific life history traits over time and among populations together with changes in population size can explain X:A polymorphism ratios observed across human populations. A) Changes in population sizes in six human populations inferred by pairwise MSMC (44). Split times among populations were determined visually and are marked by stars: blue for (YRI, non-African populations), black for ((CEU, TSI), (CHB, JPT), MXL), purple for (CEU, TSI), and red for (CHB, JPT). B) Comparison of estimated normalized, polymorphism X:A ratios with those predicated under the historical changes in population size shown in A, and assuming: i) no life history effects (black); ii) sex-specific life history parameters that vary among the demarked intervals and were chosen to best fit the estimates (red); iii) the best fit further allowing sex-specific life history parameters to vary among populations after they split (blue). See text and SI Section 3 for details.

### Polymorphism ratios in six populations

Next, we examine whether variation in life history can explain the polymorphism ratios observed in all six populations jointly. For comparison, we first consider the model without sex-specific life history, which expectedly yields a poor fit (Fig. 7B). Next, we allow for sex-specific life history parameters (within the ranges detailed above) and let them vary among the intervals defined by the approximate split times among populations (Fig. 7A). In particular, we seek the parameter values that minimize a weighted squared distance between predicted and estimated polymorphism ratios (see SI Section 3). Allowing sex-specific life history parameters to vary over time but not among populations substantially improved the fit, but fails to account for some features, e.g., the ratio in YRI (Fig. 7B). Further allowing sex-specific life history parameters to differ after populations split from one another, we are able to closely match the point estimates for all six populations (with mean distance < 0.11 SEM averaged over the observed estimates; Fig. 7B).

### Life history traits during human evolution

Our results illustrate that historical changes in sex-specific life history traits and in population size can explain the X:A polymorphism ratios in extant human populations. Our analysis relied on somewhat arbitrary decisions to fit few extant polymorphism ratios using many ‘historical’ life history parameters, i.e., about possible parameter ranges, the time intervals in which they could vary, and the distance between predictions and estimates that was minimized. Alternative decisions would doubtless result in other sets of parameters that match the estimates of polymorphism ratios to a similar degree (accounting for uncertainty). The specific set of values we found (Fig. S6) should therefore be treated as one of many possibilities; narrowing these sets down will require bringing to bear richer summaries of the data (see Discussion). Nonetheless, our results suggest a few conclusions. The first is that ancestral human populations were highly polygynous, as explaining the polymorphism ratios in YRI would be difficult otherwise. Second, they indicate that non-African populations likely experienced a substantial reduction in polygyny and/or increase in male-biased generation times around the OoA bottleneck, helping to explain the large reduction in polymorphism ratios in non-African populations. Third, we find that, quite surprisingly, fairly recent changes in sex-specific life history have had a substantial impact on extant diversity levels, and in particular can account for the reduction in ratios between European and Asian populations.

## Discussion

Life history traits, and generation times in particular, affect X:A polymorphism ratios in multiple ways, and these effects can be surprisingly strong. In particular, we have shown that in humans, higher generation times in males than in females substantially decreases the mutation rate and increases coalescence rates on the X relative to autosomes. They also substantially enhance the reduction in the X:A ratio due to bottlenecks (or alternatively increase the ratio due to population growth), both by accelerating the response time in generations and by increasing the number of generations per unit time on the X relative to autosomes. These generation times effects compound those of higher reproductive variance in males (i.e., polygyny) that were explored by previous studies (1, 4, 8, 11, 21, 25). Higher male variances decrease coalescence rates on X relative to autosomes and dampen the effects of changes in population sizes on X:A diversity ratios. As we show, considered jointly, these effects can explain observed X:A polymorphism ratios across human populations.

While our results have clear implications about the values of life history traits in recent human evolution, our ability to draw quantitative conclusions is limited by remaining gaps in our knowledge about demographic history. Current demographic inferences assume that the autosomal generation time and mutation rate were constant, whereas both have doubtless changed over time. Ignoring such changes introduces errors in estimates of effective population sizes and in their assignment to past dates (i.e., in years). Accounting for these errors is unlikely to change our qualitative conclusions, but would likely affect the life history parameter estimates. The same is true of demographic complications that we did not consider, including historical migration/admixture among populations (45) and ancient introgression (46, 47), and more speculative sex biases in these processes (8, 9, 48).

Our analysis further relies on pedigree-based estimates of mutation rates in contemporary humans in order to model mutational effects on X:A polymorphism ratios. We used these estimates to infer X:A genealogical ratios and to relate models of historical life history trait values with extant X:A polymorphism ratios. In so doing, we assumed that mutation rates on the X are well approximated by the averages over rates in males and females, i.e., that 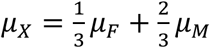 (because pedigree-based estimates rely on autosomal mutations). Although this assumption is inexact, given the evidence for X specific modifiers of mutation rates, the observed effects are fairly subtle (49). In the future, larger pedigree studies in humans, with a sufficient number of mutations on the X, should allow direct estimation of mutation rates on the X. Our approach also assumed that male mutation bias and its dependence on generation times observed today hold for the entire period over which extant neutral diversity in human arose, e.g., over the past ~1.5 MY (43). The evidence regarding evolutionary change in male mutation bias is contradictory. Lineage-specific, divergence-based estimates of *α* in great apes are extremely variable, with estimates of 2.81 for humans, 5.13 for chimpanzees, 1.15 for gorillas and 1.96 for orangutans (34) (some but probably not most of this variation could be due to changing sex-ratios of generation times (36)). In contrast, pedigree-based estimates of *α* in extant species spanning a much greater phylogenetic range, i.e., mammals, appear to be stable (albeit with large confidence intervals), and are in fact consistent with the estimates in humans ((50–55); Felix Wu and Molly Przeworski, personal communication). Larger pedigree-based studies in other catarrhine species will likely resolve this apparent conflict and inform the plausibility of our assumption. More generally, we note that pedigree-based estimates of the autosomal mutation rate have triggered a wholesale revision of the chronology of human evolution obtained from genetic data (43). Similarly, our results call for a revision of human X:A polymorphism ratios in light of pedigree based estimates of the male mutation bias.

Novel insights about mutation may also facilitate direct inferences about historical changes in life history traits. Such inferences could rely on the fact that different kinds of mutations have distinct dependencies on male and female generation times (32) but share the same genealogies. It may therefore be possible to infer male and female generation times from the ratios of different kinds of mutations of the same age on X and autosome linked genealogies. It might also be possible to extend methods like MSMC to utilize data about different kinds of mutations on the X and autosomes jointly, in order to infer historical changes in both generation times and effective population sizes, and possibly even sex-dependent migration between populations.

While we focused our analysis on humans, for which there are more data, there is every reason to think that life history substantially affected X:A polymorphism ratios in other species as well. Notably, sex differences in life history traits and changes in population size, as well as extensive variation in these factors among populations and closely related species, are pervasive in many taxa (e.g., among vertebrates (56)). It almost necessarily follows that the life history and particularly the generation time effects that we describe would have affected their X:A polymorphism ratios.

## Acknowledgements

We thank I. Agarwal, P. Moorjani and M. Przeworski for many helpful discussions and comments on the manuscript. We also thank the editor and three anonymous reviewers for many helpful comments on an earlier version of this manuscript.

