## Supplemental Information for "Changes in life history and population size can explain relative neutral diversity levels on X and autosomes in extant human populations"

Figure S6. Life history parameters that maximize the fit between predictions and observations. 17

### 1. Pedigree-based estimates of male mutation bias ($\alpha$)

We rely on the largest pedigree-based study of de-novo mutations to date, Jónnson et al. (1), to evaluate the dependence of male mutation bias, $\alpha$, on male and female generation times. Specifically, we use the linear fits for the dependence of mutation rates on generation times in each sex from Gao et al. (2), to obtain

$$\alpha(G_M, G_F) = \frac{\mu_M(G_M)}{\mu_F(G_F)} \cong \frac{1.41 \cdot G_M + 5.5}{0.39 \cdot G_F + 2.04}. \quad \text{(S1)}$$

Our results are insensitive to the particular choice of linear fit (i.e., from (1) or (2)), although we note that both fits are limited by high sampling error. Moreover, neither fit corresponds exactly to the same genomic regions from which we inferred X:A ratios and both include CpG mutations, which have been excluded from our analysis. Unfortunately, we cannot account for these effects, because the data to so was not provided by Jónnson et al. (1).

### 2. Estimating polymorphism ratios in human populations

#### 2.1. Data sources and rudimentary analysis

**Polymorphism data.** We used 1000 Genomes Project (phase 3) data (3). In order to avoid differences in mean coverage between X and autosomes, we restricted the analysis to females (Table S1). We kept only high quality biallelic SNP and monomorphic sites that passed the 1000 Genomes Project "strict" filter. In addition to the 1000 Genomes Project call mask, we removed assembly gaps, centromeres, telomeres and 2Mb from the 5' and 3' end of each chromosome. We filtered simple repeats, low complexity and segmental duplications from repeatmasker files downloaded from UCSC (4). We also removed CpG sites throughout the genome as well as all sites within CpG islands and the pseudoautosomal region of the X.

To select putatively neutral sites, we first removed all sites with phastCons 100-vertebrate alignment scores greater than 0 (sites without scores were also removed) (5). We also removed all exonic sites and sites 1kb upstream, 500 bp downstream, and the first and last 200 bp of introns corresponding to our set of exons (see details on exonic coordinates below). Since we estimated divergence at neutral sites by aligning to rhesus macaque (rheMac3), we also removed sites that did not align to the macaque genome. After applying these filters, we obtained 715,834,113 putatively neutral sites on autosomes and 23,304,829 on the X. For the analysis described in Section S2.3, we also estimated divergence from an alignment to orangutan (ponAbe2), in which case we applied the same filters but removed sites that did not align to the orangutan rather than macaque genome. With the alignment with orangutan, we obtained 728,753,605 putatively neutral sites on autosomes, and 23,824,929 on the X

**Polymorphism levels.** At each putatively neutral site, we calculated heterozygosity ($\pi$) for each population sample using the formula $2PQ/N(N-1)$, where $P$ and $Q$ are the numbers of chromosomes in the sample carrying each allele, and $N = P + Q$.

**Divergence levels.** Divergence from macaque or orangutan for putatively neutral sites in a bin (see definitions of bins below) was estimated by dividing the number of sites at which human and macaque or orangutan carried distinct alleles by the total number of sites.

We used divergence levels as a proxy for substitution rates, resulting in underestimates of substitution rates due to multiple hits. Average divergence levels between human and macaque in our filtered set of sites (which excludes CpG sites) are ~6.5% on autosomes and ~5.7% on the X. Assuming a simple, Jukes-Cantor mutational model, these values suggest that we underestimated the substitution rate by ~7.9% on autosomes and by ~7.1% on the X (i.e., the estimated substitution rates should be ~7.1% and ~6.1%, respectively), and that accounting for multiple hits would increase our estimates of the X:A polymorphism ratios by a multiplicative factor of 1.01. For divergence between human and orangutan the effect is even smaller. While this is a rough approximation, it indicates that the effect of multiple hits is tiny relative to our confidence intervals, which is why we chose not to correct for it. We chose to exclude CpG sites because divergence rates at these sites are a poor proxy for substitution rates (due to multiple hits and, more so, to their mutation rate dropping after they mutate).

**Exon coordinates.** We used exon coordinates to filter sites and to bin neutral sites by genetic distance to the nearest exon. To construct a set of exons, we downloaded UCSC hg19 knownGene files (4) and collected exons for all protein coding genes. For genes with multiple splice variants, we kept the set of exons in the longest isoform. In the rare cases of two overlapping gene predictions, we retained the gene with the longer exon sequence.

**Genetic Map.** To obtain genetic distances to exons, we used the Hinch et al. genetic map, which was constructed based on switches in ancestry tracts in African-Americans (6). Among high-resolution genetic maps in humans, this one is likely the least confounded by polymorphism levels. We rescaled the genetic map for the X by ⅔, to obtain sex-averaged rates (Hinch et al. report rates in females for the X, as they normalize by total genetic lengths obtained from HapMap). The genetic distance between a neutral site and an exon was defined as the distance to the exon's midpoint.

**Confidence Intervals.** Confidence intervals on all our estimates were obtained using bootstrapping, in which we re-sampled 1 MB windows with replacement, excluding windows without any putatively neutral sites.

### 2.2. Estimating X:A ratios in the absence of selection at linked sites

**Estimating ratios far from genes.** The standard method to estimate $\pi/D$ while minimizing effects of selection at linked sites is to measure polymorphism and divergence far from genes (7). To this end, we binned putatively neutral sites based on genetic distance (in cM) from exons. We used 102 bins in total, starting with smaller bins of [0, 1e-04), [1e-04, 1e-03), [1e-03, 1e-02), to capture the effects of selection (8), and then continued with intervals of 0.01 (i.e., [0.01, 0.02), [0.02, 0.03), etc.) up to [0.99, 1.00); the final bin was [1.00, 10) and captured the relatively few remaining sites at distances ≥ 1 cM (Fig. S2A). We calculated the average polymorphism and divergence levels in each bin, and estimated the ratio by $\bar{\pi}/\bar{D}$. To estimate the value of $\pi/D$ at genetic distances greater than a specified threshold, we averaged our estimates of $\pi/D$ across bins above the threshold, weighting bins by the number of sites in them. We then divided our estimates for the X and autosomes in order to obtain an estimate of the X:A ratio corresponding to a given threshold.

The choice of threshold reflects a trade-off between accuracy and precision. If the threshold is too low then the estimate might be affected by selection (Fig. S2B), and if it is too high then the estimate would rely on small sample sizes and have large confidence intervals (Fig. S2A, C and D). Using a threshold of 0.2 cM suggested by previous work (9) restricts the analysis to 24% of the sites on autosomes and 17.6% on the X (Fig. S2A). The McVicker et al. $B$-maps suggest that diversity levels at this distance (and even much farther) are still affected by selection at linked sites, but that with a threshold of 0.2 cM, the effects on the X and autosomes approximately cancel out (with average McVicker's $B$ of 0.939 on autosomes and 0.941 on the X; Fig. S2B). We therefore used this threshold for our estimates (Fig. S5, Table S2).

**Estimating ratios using McVicker's $B$-maps.** An alternative approach, which should improve precision, is to estimate neutral values of $\pi/D$ without imposing a threshold on the distance from genes, but correct for the effect of selection at linked sites using the McVicker et al. $B$-maps (10). To this end, we binned putative neutral sites by their estimated $B$ value (using the bins in which $B$ values were reported (10)), and estimated the average values of $\pi$ and $D$ for each bin. We then calculated the average value of $\pi/BD$ across all bins, weighting bins by the number of sites in them. In theory, estimates of $\pi/BD$ should not depend on $B$, as the division by $B$ should remove the effects of linked selection, and the division by $D$ should corrects for correlations between $B$ values and local mutation rates.

In practice, we find indications for biases in estimates of $\pi/BD$ at the lower and upper range of $B$-values (Fig. S3). Estimates of $\pi/BD$ for $B$ values near 0 are much greater than in the rest of the range (Fig. S3C), plausibly due to the presence of segregating sites near regions that were incorrectly annotated as highly conserved in the inference of $B$ values. Also, on the X, estimates of $\pi/BD$ in this range exhibit large fluctuations, plausibly due to small sample sizes (Fig. S3B). At the high end of the range, e.g., $B > 0.95$, $\pi/D$ appears to increase super-linearly (Fig. S3B), perhaps due to increased introgression from archaic humans in regions that are both highly recombining and functionally depleted (11, 12). Given these observations, we estimated X:A ratios based on averaging $\bar{\pi}/B\bar{D}$ over bins with $B$ values between 0.6 and 0.9 on the X and autosomes. This range includes ~44% of putatively neutral sites on autosomes and ~28% on the X (Fig. S3). While the thresholds of 0.6 and 0.9 are somewhat arbitrary, our estimates for X:A ratios are fairly insensitive to these thresholds, as long as sites with low $B$ values are removed (Fig. S4). Our estimates for different populations with 95% confidence intervals based on bootstrapping are shown in Fig. S5 and Table S2. These estimates are consistent with those based on regions far from exons (Fig. S5), but their confidence intervals are smaller as they rely on more sites (Figs. S2A and S3A).

#### 2.3. Estimating X:A divergence ratios at putatively neutral sites

In order to compare divergence- and pedigree-based estimates of the male mutation bias, $\alpha$, we measured ratios of X:A divergence between human-macaque and human-orangutan. We then translated these ratios into estimates of $\alpha$ using Miyata's formula (13), thus ignoring potential biases from sex-specific generation times (14) and lower ancestral polymorphism levels on the X (13). Both of the divergence-based estimates are substantially lower than estimates based on pedigree-studies in contemporary humans (Table S3 and Fig. 2). We further examined whether the divergence-based estimates of male mutation bias are substantially different in the regions that we used to infer X:A polymorphism ratios (SI Section 2.2) and genome-wide (e.g., due to difference in base composition). Comparing estimates from putatively-neutral sites genome-wide, far from exons, and in sites where $0.6 \leq B \leq 0.9$, we find negligible differences in base composition and in estimates of $\alpha$ (Table S3).

### 3. Fitting life history models to observed X:A ratios

In the main text (Figs. 7), we relied on published inferences of  historical changes in population sizes (15), and considered three models for historical values of life history traits:

- Model 1: Demography alone (black circles in Fig. 7B). We assumed that generation times and reproductive variances were identical between the sexes.
- Model 2: Identical sex-specific life histories across populations (cyan circles in Fig. 7B). We allowed $G_M/G_F$ and $(V_M + 2)/(V_F + 2)$ to vary in time and take different values in each of the time intervals defined by population split times (see below).
- Model 3: Population-specific life-histories (yellow circles in Fig. 7B). We allowed sex ratios of generation times and reproductive variances to vary among populations after they split.

In all cases, we further assumed a constant autosomal generation time of 30 years, in accordance with the assumption of the demographic inferences we relied on (15).

We defined split times visually based on where the estimated population sizes in different populations diverged (marked by '*' in Fig. 7A). Specifically, these times were chosen to occur at the ends of time intervals used by the pairwise-MSMC on which we relied (15). We denote split times as follows: $T_4$ for the split between YRI and the non-African populations, $T_3$ for the split between MXL and the other non-African populations, $T_2$ for the split between (CEU, TSI) and (CHB, JPT), and $T_1$ for the split between CHB and JPT. The split times: $0 < T_1 < T_2 < T_3 < T_4 < \infty$, define five time intervals in which life history ratios take specific values. We note that our qualitative results should be insensitive to the precise choice of split times.

Given the time intervals and corresponding values of life history ratios, the predicted polymorphism ratio was calculated using the recursions given in Amster & Sella (2019). Specifically, for each time interval: i) the effective population sizes for the X was calculated using Eq. S128 in SI Section 4 of (16); ii) the number of generations for the X and autosomes were calculated using Eq. S129 in SI Section 4 of (16); iii) the mutation rates per generation on the X and autosomes were calculated using the linear fits from Gao et al. (2) (see Section 1), given $G_M$ and $G_F$ (the generation times follow from the ratio $G_M/G_F$ in an interval, given that the autosomal generation time $G_A = \frac{1}{2}G_M + \frac{1}{2}G_F = 30$ years). Given these quantities for each time interval, the predicted X:A polymorphism ratio was calculated using Eqs. S131 and S135-S137 in SI Section 4 of (2). To compare this ratio with the corresponding divergence-normalized estimate, we divided the polymorphism ratio by the ratio of divergence between human and macaque on X versus autosomes, $\overline{D}_X/\overline{D}_A = 0.89$ (Table S3).

We found the life history ratios that maximize the fit between predicted and observed ratios under models 2 and 3 using constrained optimization (using the Python function *scipy.optimize.minimize*). In each time interval, $G_M/G_F$ was allowed to take values between 0.9 and 1.4, and $(V_M + 2)/(V_F + 2)$ was allowed to take values between 1 and 2.5. The optimization was set to minimize

$$\Delta^2 = \sum_p w_p \cdot \left(\frac{PR_p - ER_p}{SEM_p}\right)^2, \quad \text{(S2)}$$

where $PR_p$ is the predicted ratio, $ER_p$ is the estimated ratio, $SEM_p$ is the SEM obtained by bootstrapping, and $w_p$ is the weight, associated with population $p$. Populations from different continents, and within continents, were weighted equally, i.e., the weights $w_p$ were 0.25 for YRI and MXL, and 0.125 for the other four populations. The results of models ii-iv for the reduction in ratio in CEU relative to YRI in Fig. 6 were also obtained by bound optimization.

### 4. Software

Documented versions of all the software we used are available at https://github.com/sellalab/XA_poly.

### Supporting Tables

| Population | Number of female individuals |
|---|---|
| YRI | 56 |
| MXL | 32 |
| TSI | 54 |
| CEU | 50 |
| CHB | 57 |
| JPT | 48 |

**Table S1. Number of females per population in 1000 Genomes samples.**

| Population | X:A ratio far from exons | X:A ratio where $0.6 \leq B \leq 0.9$ | X:A ratio assuming $\alpha = 4.25$ |
|---|---|---|---|
| YRI | 0.81, [0.75,0.88] | 0.82, [0.79,0.86] | 0.92, [0.88,0.97] |
| MXL | 0.68, [0.61,0.75] | 0.67, [0.63,0.72] | 0.76, [0.70,0.81] |
| TSI | 0.70, [0.62,0.78] | 0.67, [0.63,0.72] | 0.75, [0.70,0.81] |
| CEU | 0.70, [0.61,0.79] | 0.67, [0.63,0.72] | 0.75, [0.70,0.81] |
| CHB | 0.65, [0.57,0.73] | 0.63, [0.57,0.69] | 0.71, [0.64,0.78] |
| JPT | 0.64, [0.56,0.72] | 0.63, [0.57,0.69] | 0.71, [0.63,0.78] |

**Table S2. Estimates of X:A polymorphism ratios.** Including divergence-normalized ratios at putatively neutral sites: far from exons, and for which $0.6 \leq B \leq 0.9$, and genealogical ratios obtained assuming a male mutation bias of 4.25. The estimates shown here are also shown in Figs. 4 and S5. 95% CI obtained by bootstrapping are reported in brackets.

| Subset of putatively neutral sites | $\bar{D}_X/\bar{D}_A$ | $\hat{\alpha} = f^{-1}(\bar{D}_X/\bar{D}_A)$ | % GC (A) | % GC (X) |
|---|---|---|---|---|
| Genome-wide (H-M) | 0.88 | 2.13 | 40.7% | 40.4% |
| $\geq$ 0.2cM from exons (H-M) | 0.89 | 1.98 | 39.3% | 39.6% |
| $0.6 \leq B \leq 0.9$ (H-M) | 0.89 | 2.01 | 40.7% | 40.7% |
| Genome-wide (H-O) | 0.85 | 2.68 | 40.5% | 40.2% |
| $\geq$ 0.2cM from exons (H-O) | 0.84 | 2.75 | 39.1% | 39.5% |
| $0.6 \leq B \leq 0.9$ (H-O) | 0.87 | 2.30 | 40.6% | 40.6% |

**Table S3. X:A divergence ratios and corresponding $\alpha$ estimates.** We report the values obtained from human-macaque (H-M) and human-orangutan (H-O) divergence at putatively neutral sites in the two subsets of the genome used and genome-wide, alongside the GC content in these sets (see Section 2.3).

### Supporting Figures

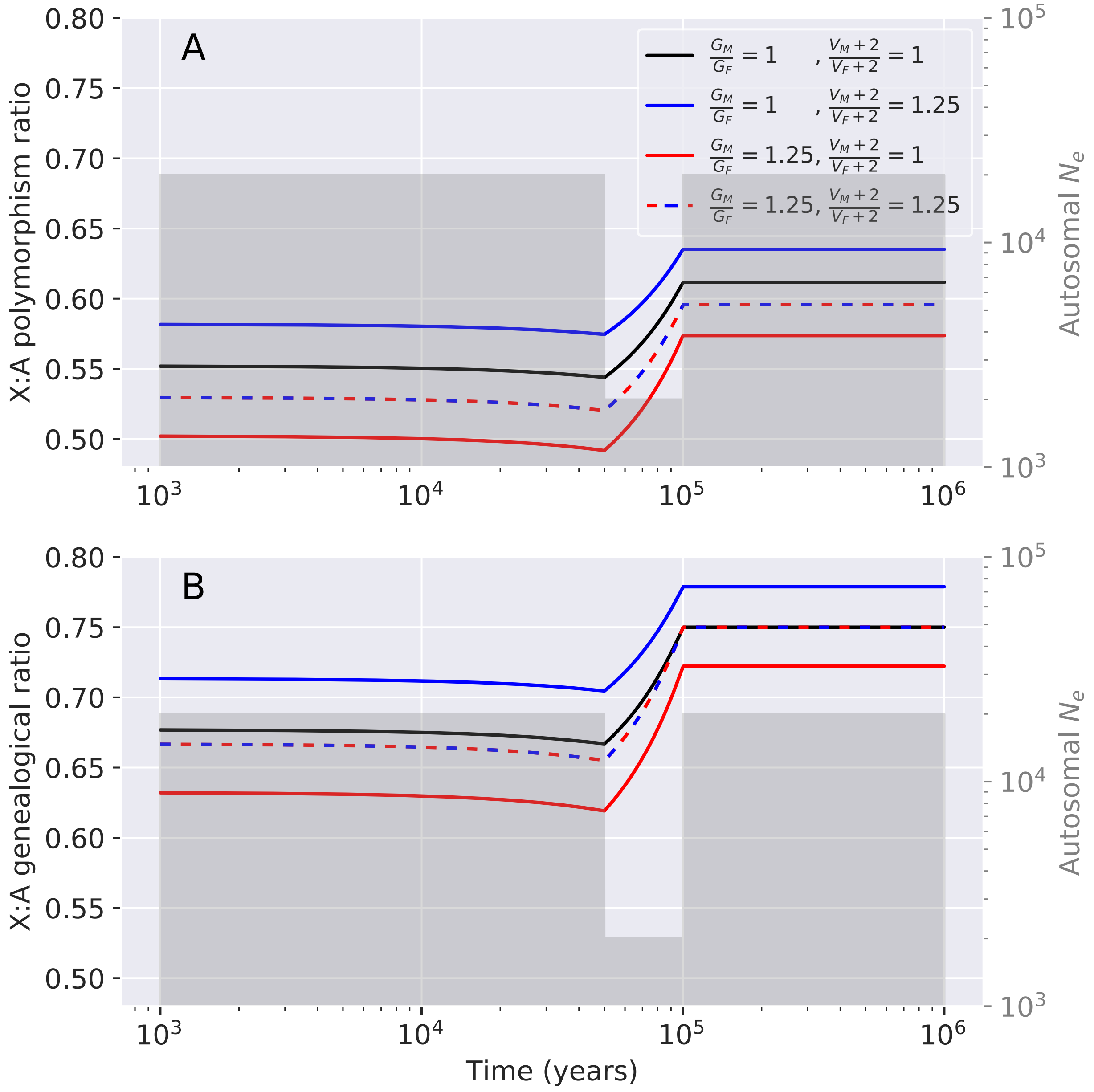

**Figure S1. Bottleneck effects on X:A polymorphism and genealogical ratios.** These figures correspond to the demographic scenario considered in Fig. 2.

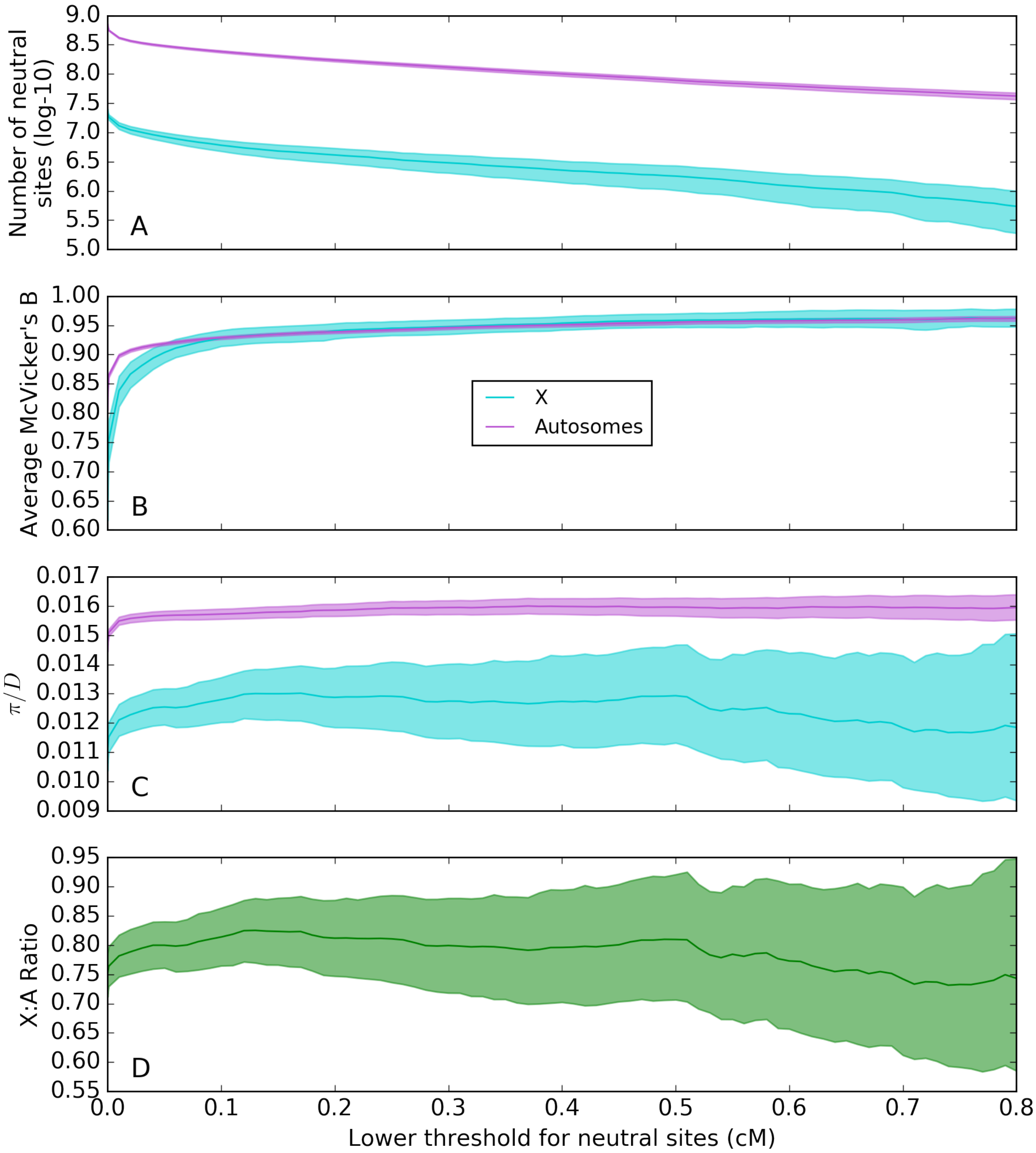

**Figure S2. Estimates of diversity levels far from exons.** Considering all sites that are beyond a certain number of cMs from the nearest exon (x-axis), we show (A) the total number of sites available for the analysis, (B) the average $B$ value, (C) the average $\pi/D$, and (D) the estimate of the X:A ratio. The shaded areas correspond to 95% confidence intervals based on bootstrapping. The estimates shown are for diversity levels in YRI and divergence between human and macaque. A similar behavior is seen in the other 1000G populations.

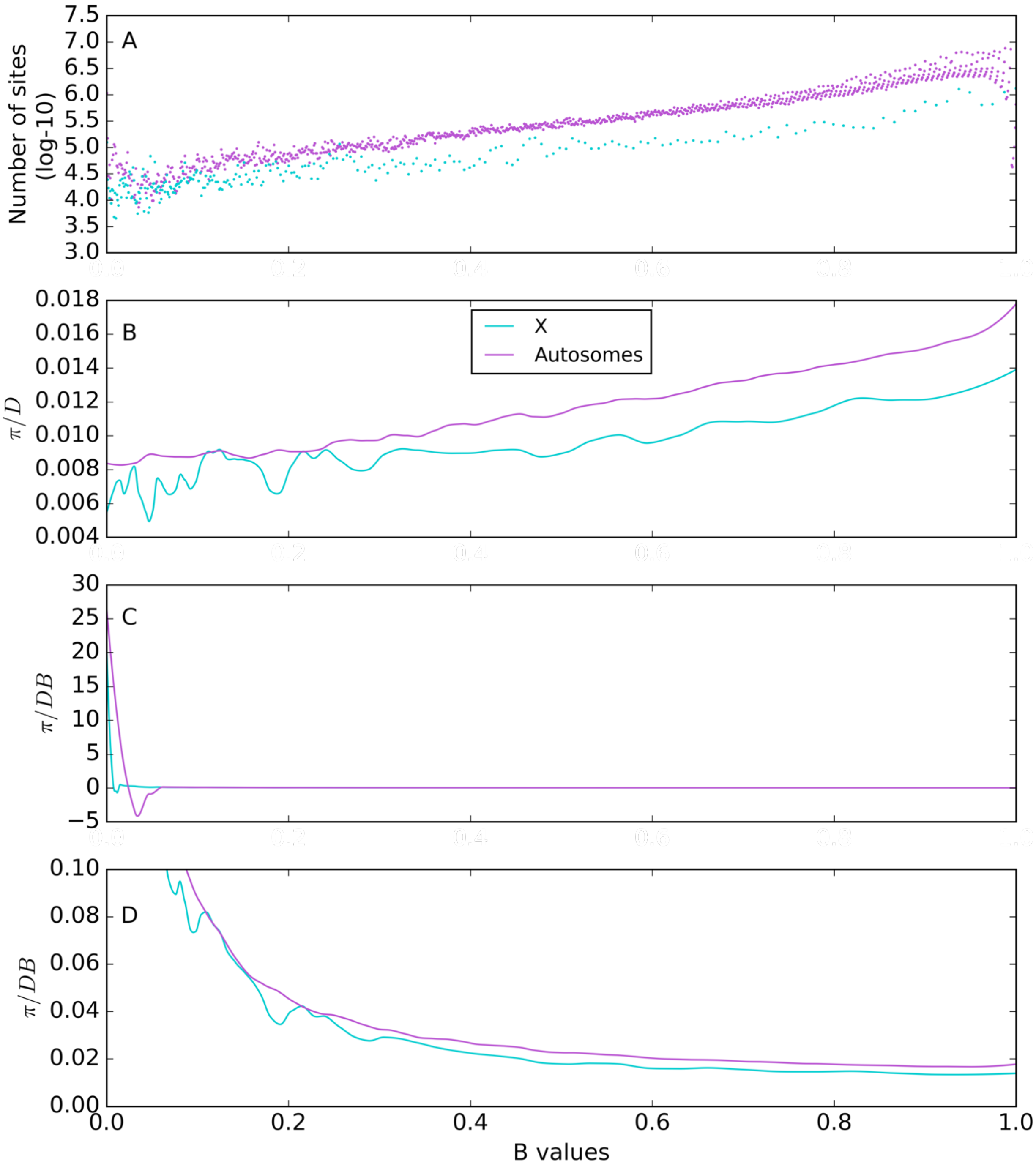

**Figure S3. Estimates of diversity levels as a function of McVicker's B.** (A) Number of sites. (B) Locally-weighted regression (loess) curve of $\pi/D$. (C) Loess curve of $\pi/BD$. Negative values are artifacts of the loess model (which is not used in obtaining our estimates). (D) Same as in C, but for $B$ values away from 0. All loess plots were generated in Python using skmisc.loess with a span of 0.1, and are weighted by the number of sites. The estimates shown are for diversity levels in YRI and divergence between human and macaque. The behavior is similar in other populations.

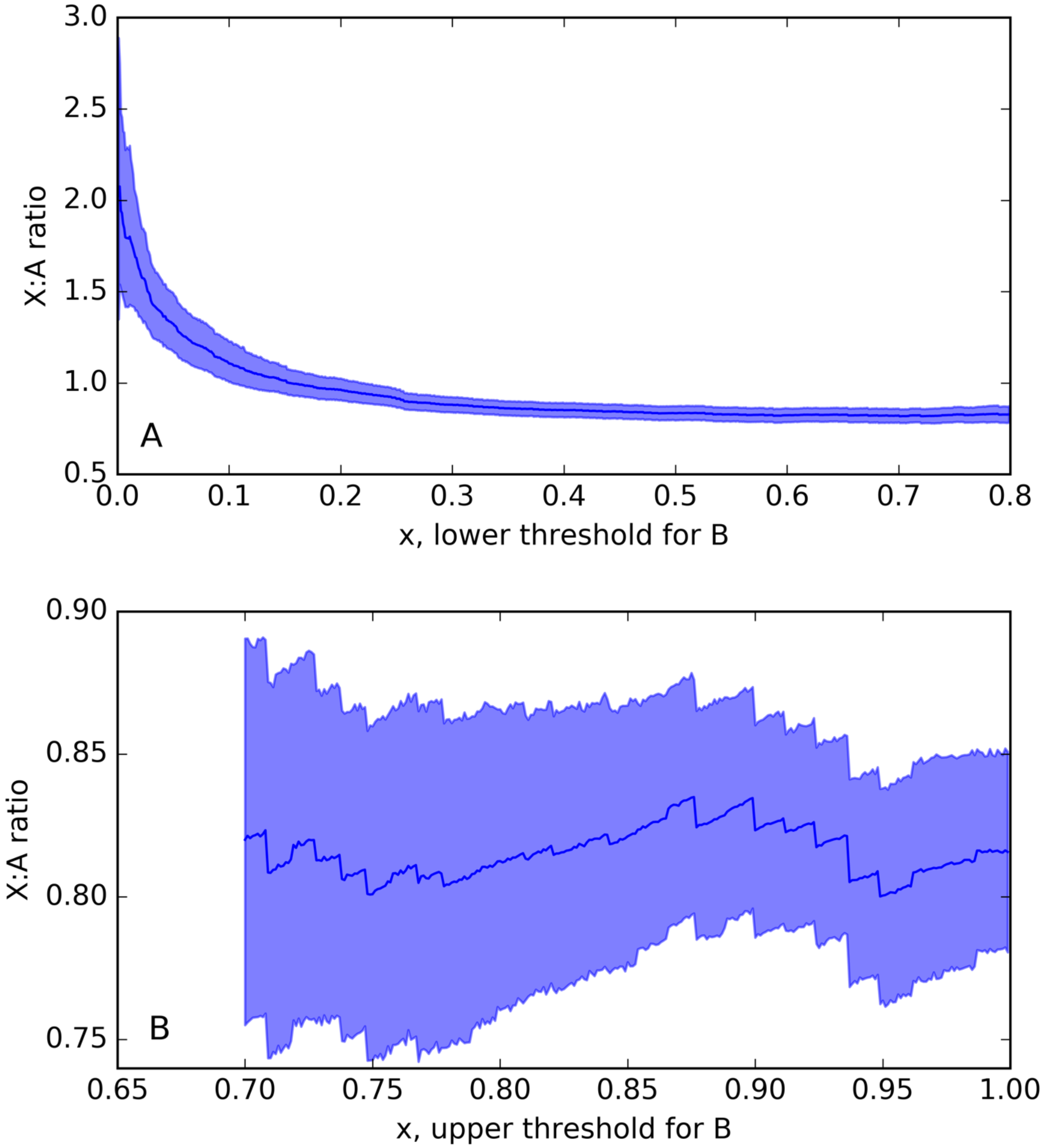

**Figure S4. Dependence of the estimated X:A ratios on the range of B values considered.** Estimates based on averaging $\pi/BD$ using sites that satisfy $x < B < 0.9$ (A), and $0.6 < B < x$ (B), where $x$ is the value shown on the x-axis. Confidence intervals are based on bootstrapping. The estimates shown are for diversity levels in YRI and divergence between human and macaque. A similar behavior is seen in the other 1000G populations.

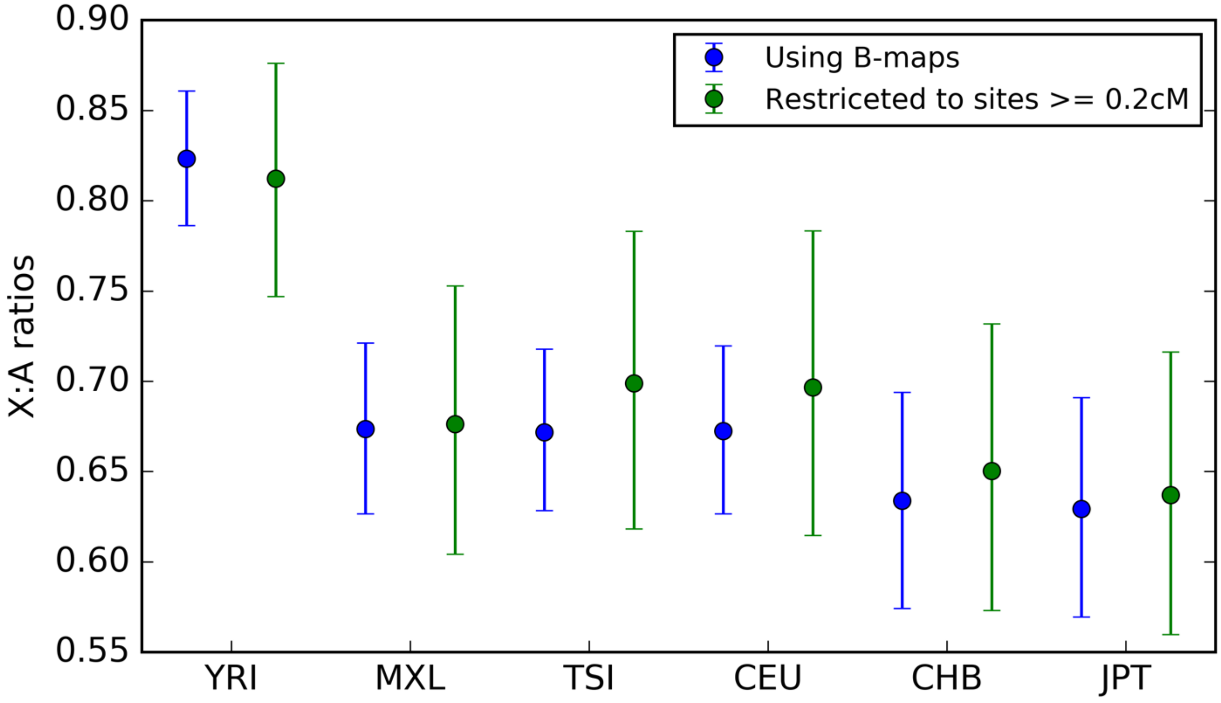

**Figure S5. Estimates of X:A ratios based on our two approaches.** Error bars represent 95% CI obtained by bootstrapping. Numerical values are reported in table S2. See Section 2.2 for details.

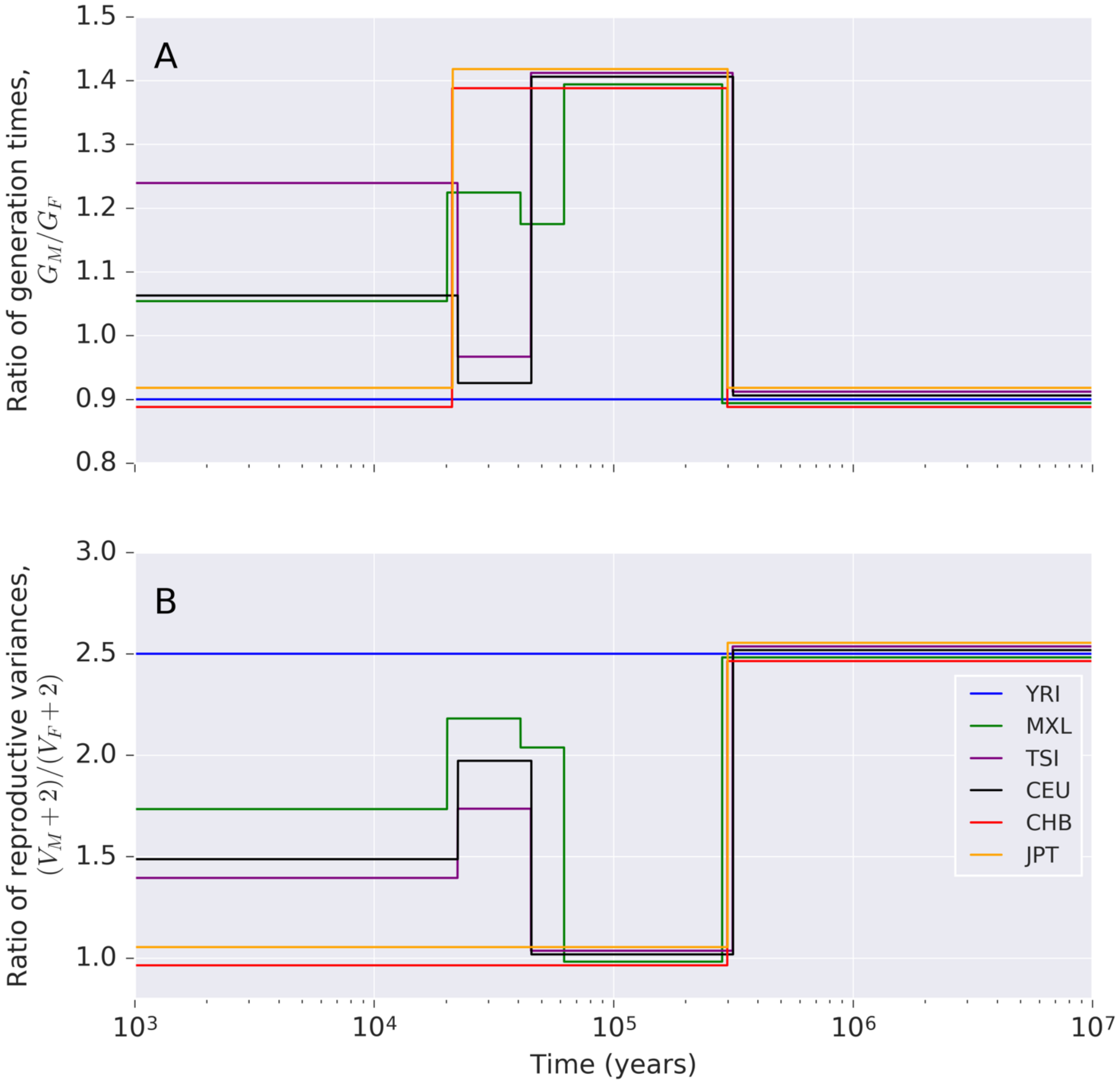

**Figure S6. Life history parameters that maximize the fit between predictions and observations.** Sex ratios of generations times (A) and reproductive variances (B) inferred by constrained optimization with the full model (corresponding to the yellow circles in Fig. 7B). In order to make identical values in different populations visually distinguishable, the values were perturbated by as much as ±0.018 (A) or ±0.054 (B). Due to the properties of constrained optimization, parameter values often 'stick' to the minimal or maximal bounds of the permitted ranges.
